# Poly ADP-ribosylation regulates *Arc* expression and promotes adaptive stress-coping

**DOI:** 10.1101/2023.09.26.558845

**Authors:** Eliyahu Dahan, Tze’ela Taub, Leah Pergamenchik, Arthur Vovk, Jade Manier, Raphael Avneri, Elad Lax

## Abstract

Rapid adaptation to stressful events is essential for survival and requires acute stress response and stress-coping strategy. Yet, the molecular mechanisms that govern this coping strategy have to be fully discovered. Poly ADP-ribosylation (PARylation) is a post-translational protein modification that plays a vital role in histone modification following neuronal activity and is required for long-term memory formation. Previous studies showed that inhibiting Parp1, the primary enzyme that catalyzes poly ADP-ribose polymers (PAR), impairs long-term memory formation. However, the target genes of Parp1-induced PARylation following acute stress are unknown. Here, we showed that the forced swim test, a well-established acute stress paradigm, induced elevated cortical PARylation and increased the expression of activity-dependent genes in mice. Systemic pharmacological Parp1 inhibition with ABT888 (Veliparib), impaired stress-coping in a second forced swim test done 24 hours later as measured by diminished immobility response. This effect was associated with reduced PARylation of Arc promoter and reduced Arc expression, 1 hour after Parp1 inhibition, suggesting Arc PARylation regulates its expression and hence successful stress-coping behavior.

## Introduction

Long-term memory formation of stressful events is an adaptive mechanism enabling the optimization of coping strategies in the future. However, under abnormal circumstances, highly stressful events and persistent reactivation of said memory can lead to undesired outcomes such as avoidance behavior and post-traumatic stress disorder (Ressler et al., 2022). The formation of long-term memories requires *de-novo* protein synthesis preceded by time-dependent transcriptional alteration (Alberini, 2008; Bisaz, Travaglia, & Alberini, 2014; Dudai & Eisenberg, 2004; Eric R. Kandel, 2001).

Poly ADP-ribosylation (PARylation) is a post-translational protein modification mostly known for its role in DNA repair. PARylation also promotes gene expression, presumably through increasing chromatin accessibility by evicting the linker histone H1 (Azad et al., 2018; Fontán-Lozano et al., 2010). The enzyme Poly (ADP-ribose) polymerase 1 (Parp1) is the most abundant member of the PARP family in mammals (Homburg et al., 2000; Ray Chaudhuri & Nussenzweig, 2017). Parp1 catalyzes the polymerization of ADP-ribose, sourced in nicotinamide adenine dinucleotide (NAD+), to yield linear or branched structured poly ADP-ribose polymers (PAR).

Previous findings by us and others indicate that Parp1 is necessary for forming and retrieving long-term memories of feeding, stress, and cocaine-seeking behaviors (Goldberg, Visochek, Giladi, Gozes, & Cohen-Armon, 2009; Hernández et al., 2009; E. Lax et al., 2017; Scobie et al., 2014). Poly ADP-ribosylation (PARylation) is a rapid and transient posttranslational modification that occurs within 10-15 minutes after depolarization and neuronal activity (Cohen-Armon et al., 2004; Homburg et al., 2000; Visochek et al., 2005). The PARylation of histone H1 regulates chromatin de-condensation and enables memory consolidation. However, it is still not clear which downstream genes are upregulated following histone PARylation. Stressful events that cause neuronal activity trigger the expression of immediate-early and activity-dependent genes required for further neuronal activation, memory formation, and neuroplasticity (Flavell & Greenberg, 2008; McEwen et al., 2015). We hypothesize that a forced swim test (FST), a highly stressful event, will initiate a fast histone PARylation alongside the mRNA expression of immediate-early and activity-dependent genes. We further speculated that systemic pharmacological inhibition of Parp1 before the FST will impair memory formation. This Parp inhibition will also reduce the mRNA expression of some known neuronal activity-related, immediate-early, and activity-dependent genes, thus allowing us to identify the mRNA expression of which of these genes are PARylation-dependent.

## Methods

### Animals

We used 8-9 weeks old C57BL/6J mice (Envigo, Israel). We included equal numbers of both male and female mice for all the experiments. Mice had ad libitum access to standard chow and drinking water and were kept at constant light-dark cycles for 12 hours each, at 22℃.

All animal experiments were approved by the Institutional Animal Care and Use Committee of Ariel University, Israel (approval No. IL-225-07-21). Institutional guidelines were followed for the proper and humane use of animals in research.

### Force Swim Test (FST)

The forced-swim test (FST) was used to induce acute stress. Mice were gently put in a glass cylinder (height: 30 cm, diameter: 10 cm) filled with water (22-33℃) with no option to escape for 6 minutes. Next, the mice were removed from the cylinder and dried in a warm, dry cage for several moments. The mice were euthanized by cervical dislocation at the following time points after FST: 15 min, 60 min, and 24h. Naïve mice not introduced to the FST were used as a control group. Cortical tissues were immediately removed, snapped frozen in liquid nitrogen, and then stored at −80℃ for later use.

### Systemic ABT-888 administration

ABT-888 (Veliparib®) is a selective PARP1 and PARP2 inhibitor with higher efficacy towards PARP1 (Thorsell et al., 2017), which crosses the brain-blood-barrier (Donawho et al., 2007). We dissolved ABT-888 hydrochloride in saline and injected 15 mg/kg of ABT888 intraperitoneally at 0.1 ml volume. The injection took place 50 min before the FST based on pharmacokinetic experiments (Donawho et al., 2007) and as we reported before (Lax et al., 2017). Equal volumes of saline were injected to control mice. For a subset of mice, twenty-four hours after the first FST, a second FST was done to assess changes in the duration of immobility as a measure of the adaptive stress response as described before (Gutièrrez-Mecinas et al., 2011; Reul, 2014; Saunderson et al., 2016; West, 1990). The experiment was video-recorded, and immobility time was manually quantified by a trained experimenter blind to the experimental conditions.

### Protein Extraction and Western Blot

Protein extraction and immunoblotting protocols were done as we described before (Lax et al., 2018). Briefly, frozen samples were lysed using RIPA buffer, loaded into 7% SDS-PAGE gel, and transferred to a nitrocellulose membrane. Membranes were first blocked with 5% non-fat milk in TBST for 1 hour and then probed with antibodies against Parp1 and PAR. Beta-actin was used as a reference protein for normalization. On completing the FST, brains were immediately removed to measure Parp1 levels and Parp1 activation (i.e., PARylation) by western blot. Detailed descriptions of the primary and secondary antibodies are provided in Supplemental Table 1. Anti-Parp1, anti-PAR, and anti-beta-actin primary antibodies were diluted 1:1000. The secondary goat anti-mouse antibody was diluted 1: 5000. To detect protein levels, membranes were exposed to an ECL solution, and images were collected using a GelDoC device (Bio-Rad). Image quantification was done with ImageJ software.

### RNA extraction, reverse transcription, and quantitative PCR (q-PCR)

We extracted RNA using an RNeasy mini kit (Qiagen) following the manufacturer’s protocol. Next, we used GoScript Reverse Transcription Mix, Oligo (dT) kit (Promega, #A2791) to reverse transcribe the RNA into cDNA following the manufacturer protocol.

The qPCR procedure was done using the Hy-syber power mix (Hylabs, Israel) and included primers for the genes listed below in supplemental Table 2. RNA expression levels were normalized to beta-actin expression levels, and values were calculated using the ΔΔCT method.

### Quantitative Chromatin Immunoprecipitation (Q-ChIP)

Mice were sacrificed, and cortices were rapidly isolated, flash-frozen, and stored at-80°C for later analysis. Samples were homogenized in 1 X PBS including 1% formaldehyde, and the homogenates were kept for 10 min at room temperature with gentle agitation. Cross-linking reactions were stopped by adding glycine (125 mM) for 10 min at room temperature with gentle agitation. Fixed chromatin samples were then homogenized in cell lysis solution (PIPES 5 mM (pH 8), KCl 85 mM, NP40 0.5%) and centrifuged for 5 min at 3000 rpm, 4°C. Pellets were resuspended in RIPA-light solution (NaCl 150 mM, SDS 0.3%, Tris-HCl 50 mM (pH 8)) and sonicated to a size range of ∼200-500bp fragments using a sonics-vibracell-vcx750 sonicator with the following parameters: 30% power, in cycles of 10 sec on/10 sec off, for a total of 36 minutes on ice. Sonicated chromatin samples were centrifuged for 2 min at full speed at 4°C. Pellets were resuspended in 1 ml of RIPA-light solution. Chromatin samples were pre-cleared with 10 µl of Magna-ChIP® protein-G magnetic beads (EMD Millipore, Cat: 16-662) pre-blocked with BSA. Next, 20 micrograms of chromatin were incubated overnight at 4°C with an anti-PAR antibody (2 micrograms, see Supplementary Table 1). Input controls were treated the same way except for not adding an anti-PAR antibody to the solution. Antibodies and chromatin were mixed with 20 µl of Magna-ChIP® protein-G magnetic beads (EMD Millipore, Cat: 16-662) for 2 hours at 4°C. The beads were then washed with low-salt wash buffer (5min; 0.1% SDS, 1% Triton-X, 2mM EDTA, 20mM Tris 150mM NaCl), then with high-salt wash buffer (5min, 0.1% SDS, 1% Triton-X, 2mM EDTA, 20mM Tris 500mM NaCl) and then with wash-solution (5min; 0.25M LiCl 1% NP-40 1% deoxycholate 1mM EDTA 10mM Tris, pH 8.0) followed by six washes with TE (5min each). Protein–DNA complexes were eluted from the beads with elution buffer (1% SDS 0.1M NaHCO3), de-cross-linked, treated with proteinase K, and purified. Purified DNA was resuspended in 100ul elution buffer. For QPCR analysis, SYBR green quantitative PCR was performed (primer sequences are listed in supplementary Table 2). To determine the relative enrichment of PAR, the 2 -ΔΔCt method was used with normalization to the input data.

### Statistics

Data are expressed as mean ±SEM unless otherwise stated. The data were analyzed by one-way ANOVA followed by Tukey’s post-hoc tests. For ABT888’s effect on immobility in the FST tests, two-way repeated measures ANOVA with time and treatment were the main factors followed by pair-wise Bonferroni-corrected t-tests. Comparisons between two groups in the qPCR and Q-ChIP experiments were made with the ΔΔCT method and one-sample two-tailed t-tests. P-values smaller than 0.05 were considered statistically significant. Possible outliers were visible in the data; therefore, we applied the Iglewicz and Hoaglin’s test for outliers under the strict criterion of z-score ≥|3.5|, which excluded no more than a single data point from each experimental group.

## Results

### FST triggers elevated cortical PARylation

Mice were exposed to the FST for 6 minutes and were sacrificed 15 min, 1 hour, or 24 hours later (Fig. 1A). PARylation was rapidly elevated 15 minutes after the FST and then returned to basal levels at 1 hour post-FST and remained at those level 24 hours later (Fig 1B; left). This elevated PARylation was followed by a slower increase in Parp1 levels, the primary enzyme that catalyzes PAR, 1-hour post-FST. Parp1 levels were reduced 24 hours later (Fig 1B; right). These findings suggest that FST triggers PARylation, in line with previous studies that showed neuronal-and neurotrophic-activity-induced PARylation (Azad et al., 2018; Homburg et al., 2000; Visochek et al., 2005).

**Figure 1:**
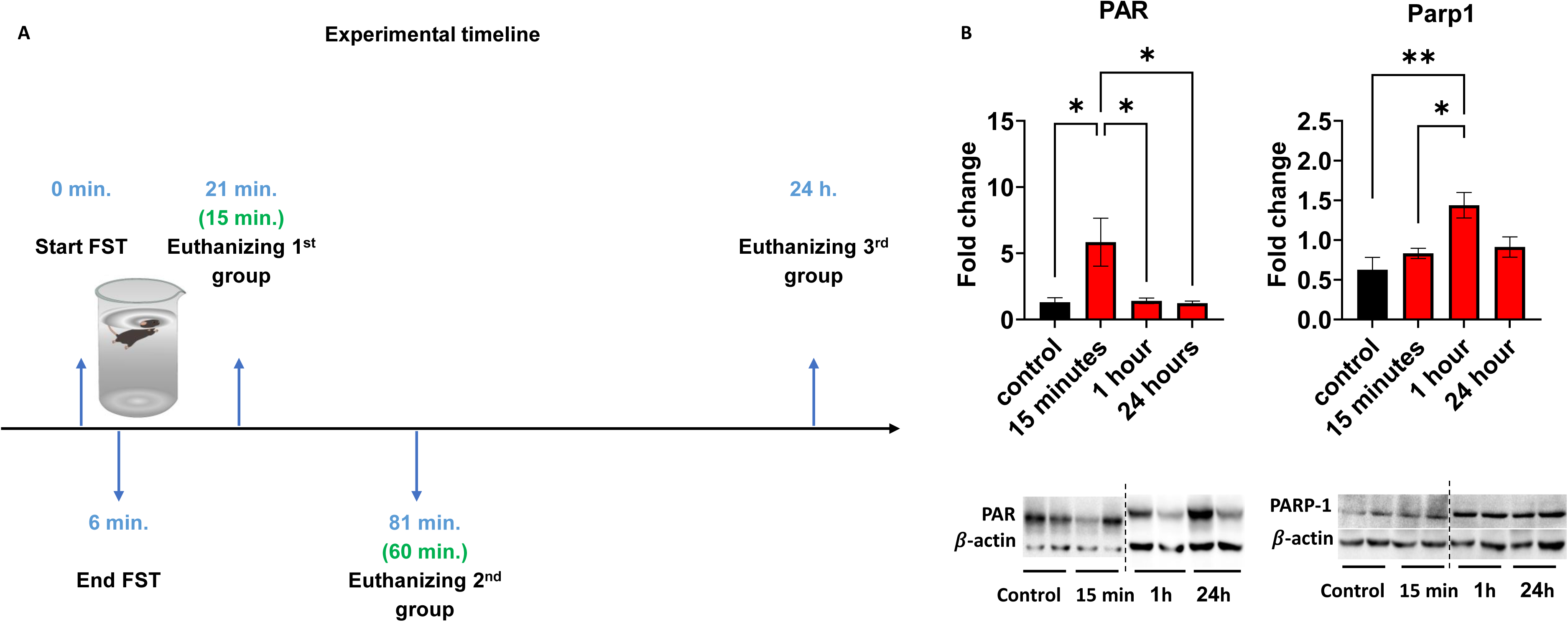
(**A**) Experimental timeline: Force Swimming Test and euthanization of the experimental groups; (**B**) Expression levels of cortical PAR and PARP1 at the time points described in (A). One-way ANOVAs for PAR levels: F (3,18) = 5.243, p=0.0089; one-way ANOVA for Parp1 levels: F (3,20) = 6.748, p=0.0025, followed by Tukey’s post-hoc test. n=5-6/group. *p<0.05, **p<0.01.

### FST induces upregulation of cortical immediate-early genes

Since FST causes extensive neuronal activation in the cortex (Connor, Kelly, & Leonard, 1997; Slattery, Neumann, & Cryan, 2011), we examined the mRNA expression levels of several immediate-early genes known to be transcribed upon neuronal activation. The mRNA expression levels were measured at the same time points as the PARylation levels (15 minutes, 1 hour, or 24 hours post-FST). We found a significant increase in Arc, c-Fos, Egr1, and Egr3 mRNA levels 15 minutes to 1 hour after the FST. The mRNA levels returned to baseline levels 24 hours later, and in the case of Egr3, mRNA levels returned to baseline levels already after 1 hour (Fig 2). Egr2, Bdnf, Nrn1, and Nr4a1 did not show a significant increase in mRNA levels, although, in all these genes, a trend toward increased mRNA expression was observed (Fig 2).

**Figure 2:**
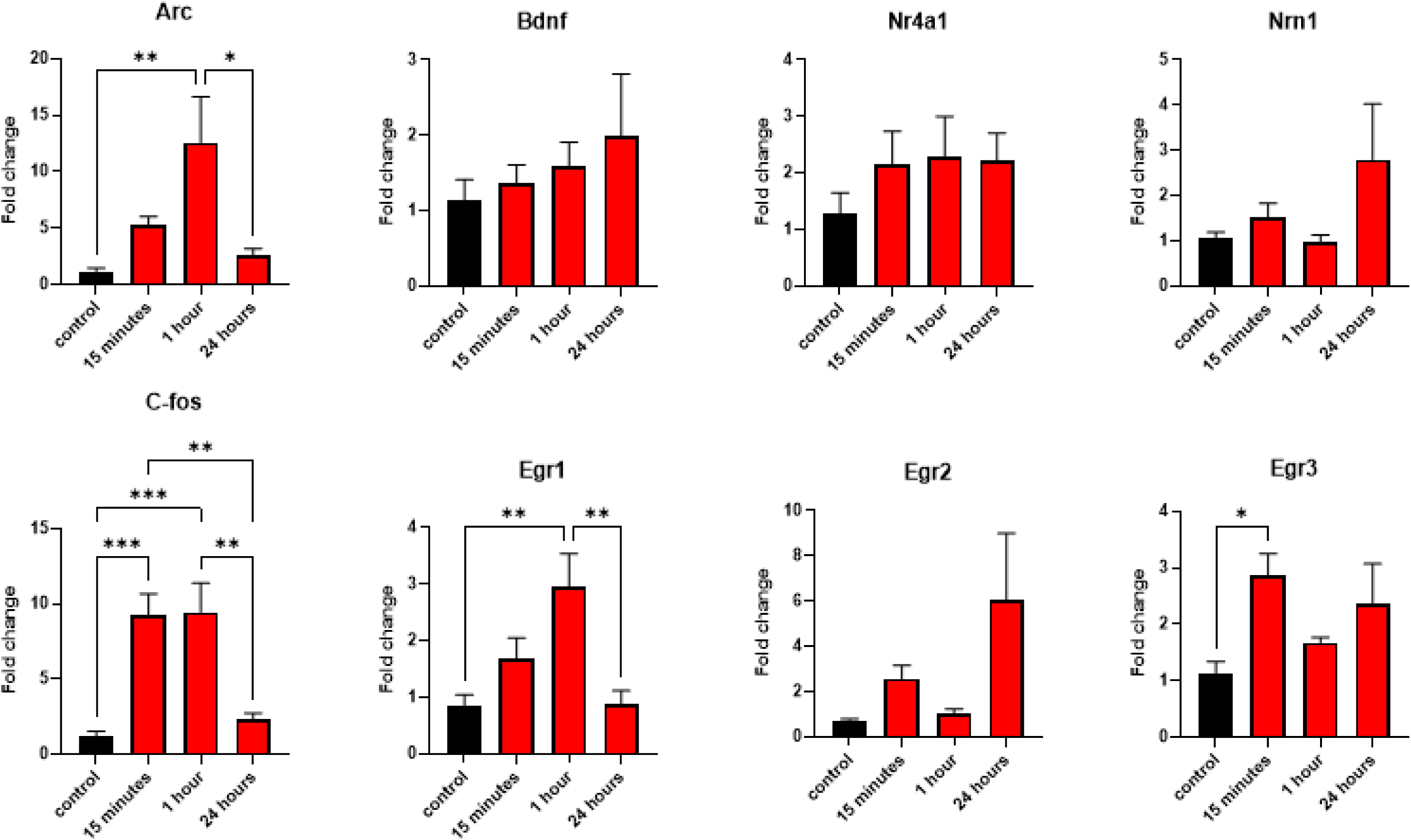
Expression levels of immediate-early and activity-dependent genes at several time points following FST. One-way ANOVAs for Arc: F (3, 20) = 5.991, p=0.0044; Bdnf: F (3, 20) = 0.5675, p=0.6428; Nr4a1: F (3, 20) = 0.7207, p=0.5513; Nrn1: F (3, 20) = 1.741, p=0.1909; C-Fos: F (3, 20) = 12.96, p<0.0001; Egr1: F (3, 20) = 7.051, p=0.0020; Egr2: F (3, 20) = 2.578, p=0.0823; Egr3: F (3, 20) = 3.400, p=0.0378. Followed by Tukey’s post-hoc test. n=6/group. *p<0.05, **p<0.01, ***p<0.001.

**Figure 3:**
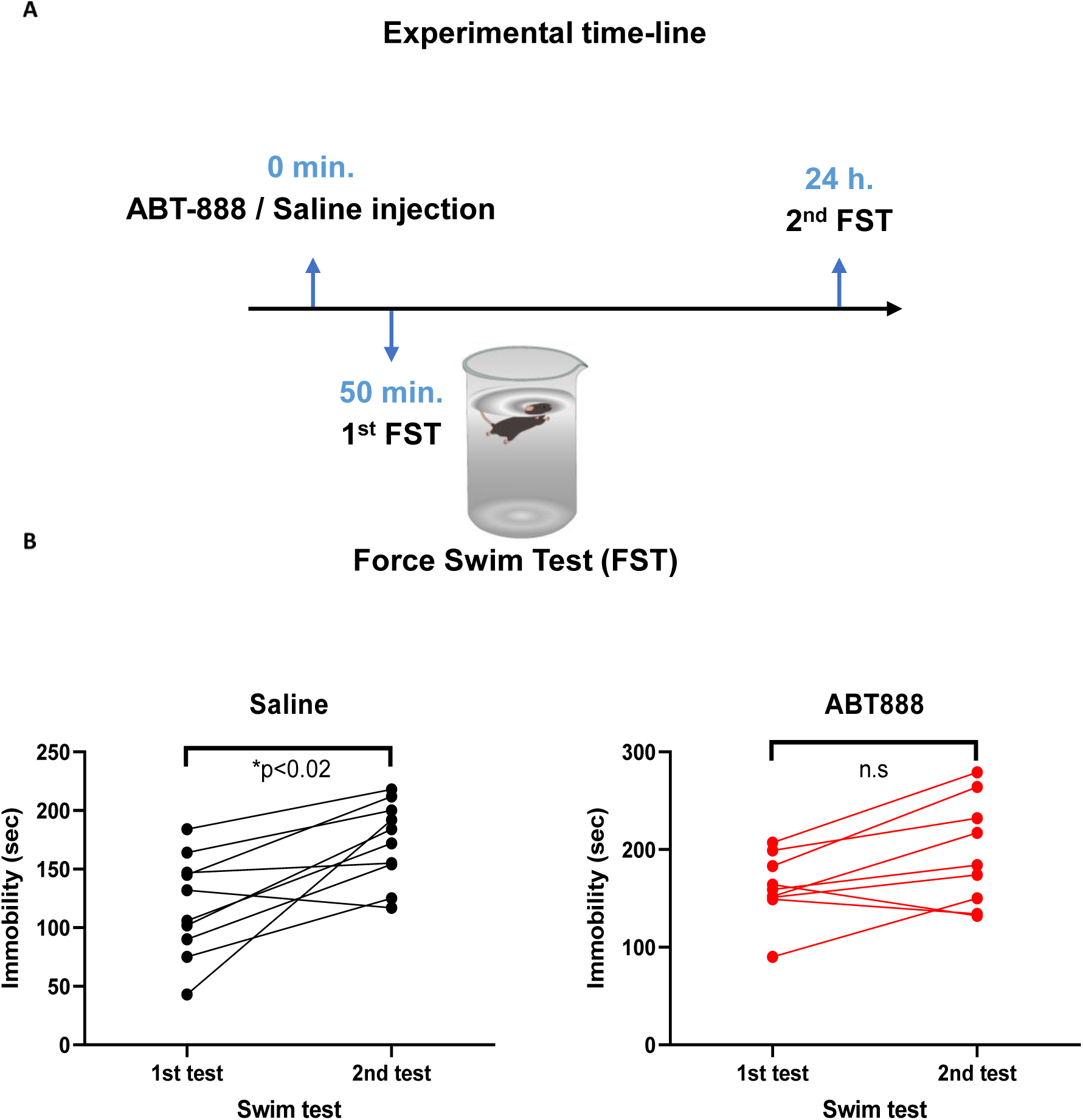
(**A**) Experimental timeline: ABT-888 / saline injection, followed by two FSTs, 24h apart. (**B**) Immobility time (in seconds) for each of the FSTs. Two-way repeated-measures ANOVA with a main effect of time F(1,17)=21.09, p=0.0003, no main effect of treatment F(1,17)=3.81, p=0.0675 and no interaction F(1,17)=1.01, p=0.3288, followed by pair-wise Bonferroni-corrected t-tests. n=10 for the saline group and n=9 for the ABT888 group. *p<0.02; n.s: no significant.

### Parp inhibition impairs stress-coping behavior in the FST

Next, we explored whether elevated PARylation induces the commonly seen shift in stress coping strategy from swimming as an active strategy into floating (i.e., immobility) as a passive strategy (Commons, Cholanians, Babb, & Ehlinger, 2017). For this aim, we injected the mice with the Parp inhibitor ABT888 (15 mg/kg IP, total volume of 0.1ml) or saline as a vehicle 50 minutes before the first FST. Twenty-four hours later, we re-exposed the mice to a second FST and recorded the immobility time during these two FST sessions (Fig 4A). We found that ABT888-injected mice showed no significant increase in immobility time, while control mice did show this shift in behavior (Fig 4B). This finding suggests that PARylation promotes the observed shift in stress coping strategy, likely through gene-expression regulation of immediate-early genes.

**Figure 4:**
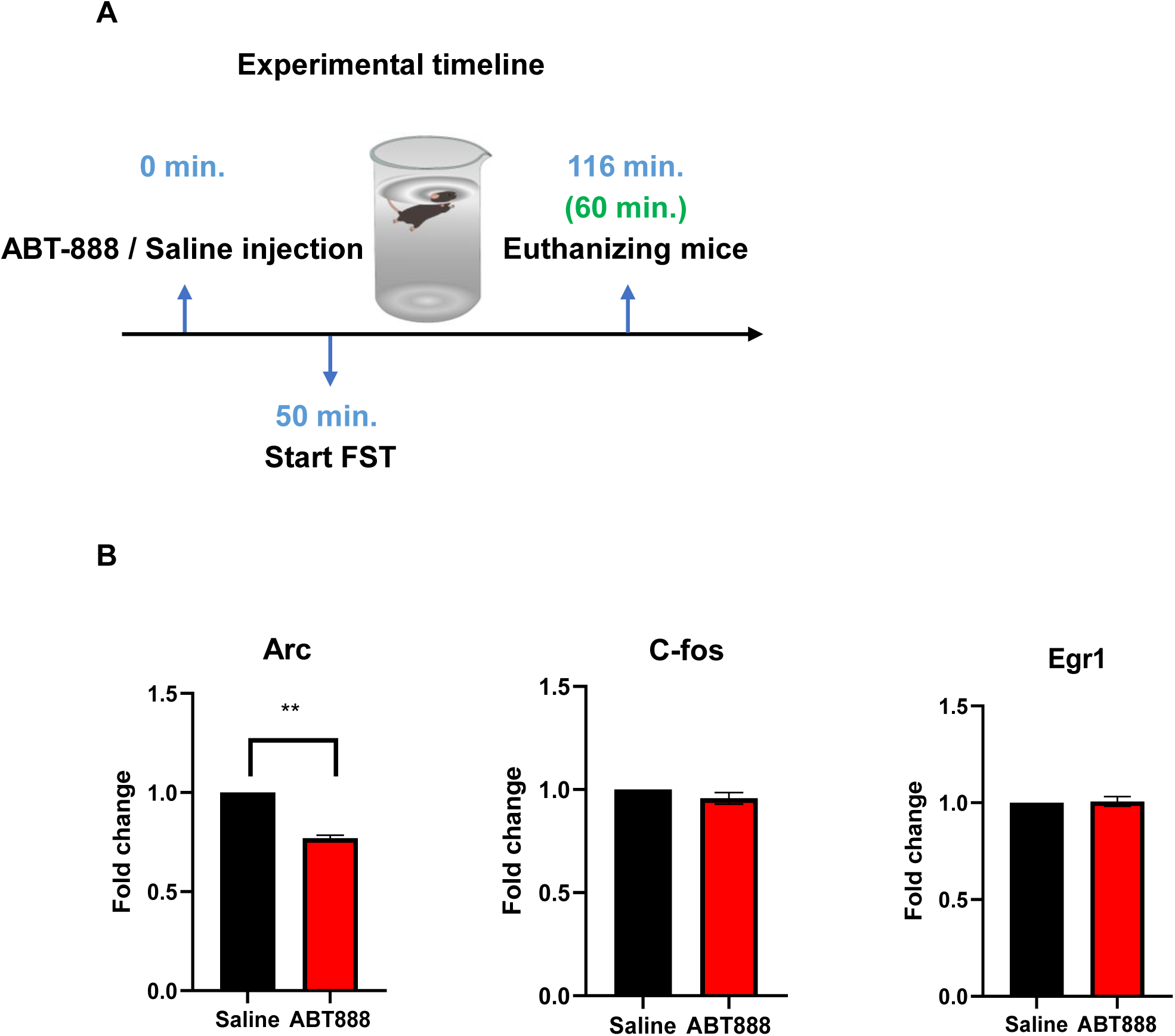
(**A**) Experimental timeline: ABT-888 / saline injection, followed by FST and mice euthanization 1 h after the end of the FST. (**B**) The relative expression level of the genes elevated 1 h after FST: Arc, C-fos, and Erg-1. T-test, **p<0.01, n=11-12 for the saline group and n=12-15 for the ABT888 group.

### Parp inhibition reduces Arc expression after FST

Based on our findings, we speculated that the FST-induced increase in mRNA expression of at least some of the immediate-early genes is due to chromatin PARylation and, hence, relaxation that enables transcription. To identify which of the immediate-early genes were regulated through PARylation, we repeated the injection of ABT888 with the same settings, introduced the mice to the FST, and one hour later, we measured mRNA levels of the immediate-early genes that were significantly elevated in the previous experiment (i.e., Arc, C-Fos, Egr1; Fig 4A). We found that Arc (but not c-fos and Egr1) mRNA expression was significantly reduced following FST due to Parp inhibition (Fig 4B). This implies that Arc is a downstream target of FST-induced PARylation.

### Parp inhibition reduces PARylation of the Arc promoter after FST

Next, to establish that PARylation directly regulates Arc mRNA expression, we injected ABT888 as before and introduced the mice to the FST. We sacrificed the mice 15 minutes or 1 hour later. We performed quantitative chromatin immunoprecipitation of PAR followed by qPCR of the Arc promoter region. We found that 15 minutes after the swim test, mice pretreated with ABT888 showed significantly lower PARylation levels on the Arc promoter compared to saline-pretreated controls (Fig 5A). This effect was not seen 1 hour after the swim test (Fig5B), in line with the transient nature of PARylation we observed before. These findings suggest that Arc expression is regulated by PARylation after the swim test to promote stress coping.

**Figure 5:**
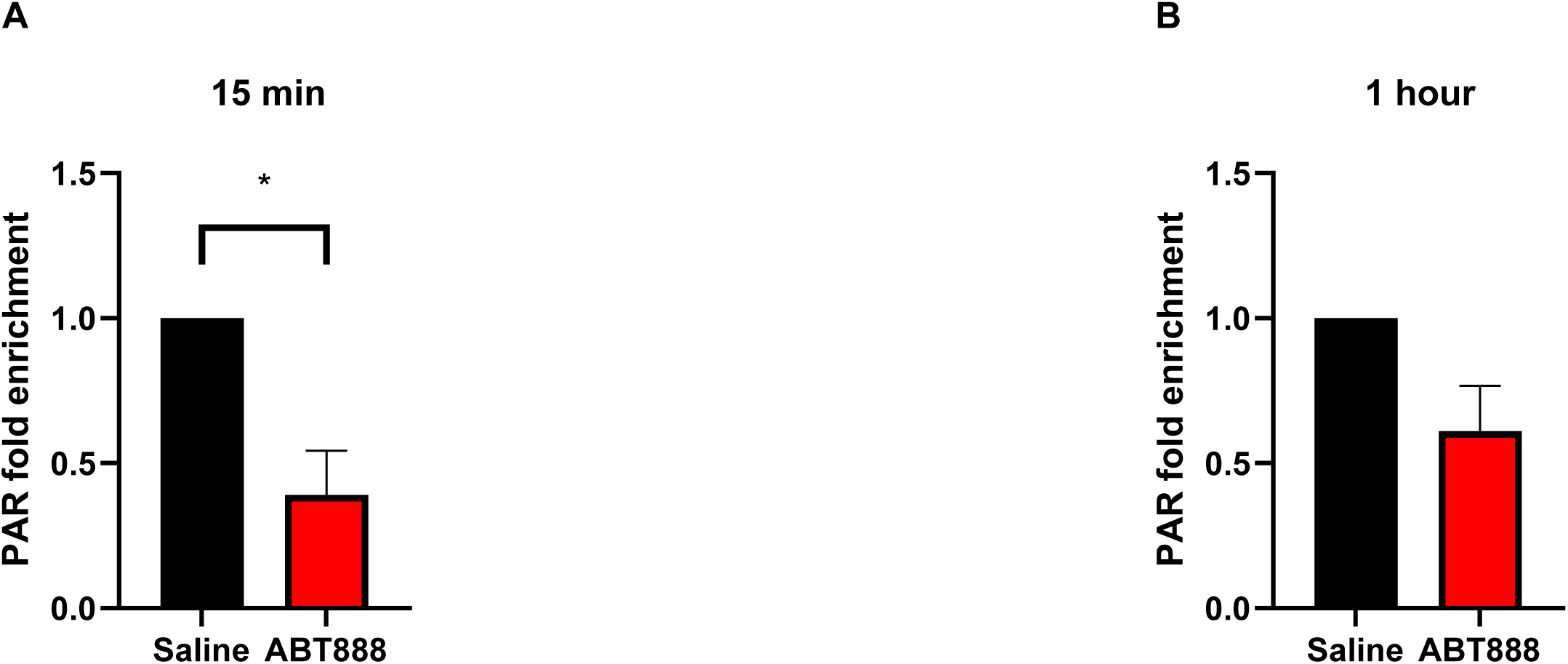
PARylation enrichment on the Arc promoter. PAR fold enrichment on the Arc promoter was significantly reduced 15 minutes **(A)** but not 1 hour **(B)** after the FST in ABT88 pretreated mice. T-test, *p<0,05, n=6 per group for the 15-minute experiment and n=3-4 per group for the 1-hour experiment.

## Discussion

Neuronal and behavioral responses to stressful events are required for proper adaptation to stress. However, the underlying molecular mechanisms that govern this response are yet to be fully explored. Here, we showed that PARylation, a necessity for long-term memory formation (Cohen-Armon et al., 2004), is induced shortly after exposure to FST and promotes adaptive stress coping.

PARylation happens shortly after neuronal activation or a neurotrophic stimulus (Homburg et al., 2000; Visochek et al., 2005; Wang et al., 2012). Parp1 catalyzes PAR chain formation on itself (i.e., auto-PARylation) and nearby histones, thus likely increasing chromatin accessibility (Azad et al., 2018; Kraus & Lis, 2003). This, in turn, enables elevated gene expression of immediate-early genes. An earlier *in-vitro* study found that high-frequency stimulation of cultured rat cerebral neurons induced PARylation-mediated expression of Arc, c-Fos, and Egr1 (Visochek et al., 2016). However, whether PARylation-mediated expression of these and other immediate-early genes occurs *in-vivo* in response to stressful stimuli was unknown. We hypothesized that (a) PARylation would be induced upon FST as part of the expected cortical neuronal response and stress adaptation; (b) that transcription of neuronal activity-dependent immediate-early genes that promotes neuronal plasticity and long-term memory formation are likely targets of this FST-induced PARylation (Cohen-Armon et al., 2004; Goldberg et al., 2009; Hernández et al., 2009; Visochek et al., 2016); and (c) that Parp inhibition will reduce the expression of these genes upon FST, resulting in reduced immobility in the second FST due to impaired adaptation response to the stress induced by the first FST.

As expected, we found that FST rapidly induced elevated cortical PARylation 15 minutes after FST, accompanied by a slower increase in Parp1, the primary enzyme that catalyzes PARylation. Further, systemic Parp1 inhibition diminished adaptive stress response in a second FST test. Gene-expression analysis of genes that were elevated after FST revealed that Parp1 inhibition reduced the expression of Arc, a known regulator of synaptic plasticity and protein-synthesis-dependent long-term potentiation (Korb & Finkbeiner, 2011; Messaoudi et al., 2007). Also, using chromatin immunoprecipitation, we demonstrated ABT888 successfully reduced PARylation of the Arc gene promoter 15 minutes after FST, suggesting PARylation has a direct regulatory role on Arc expression. These findings partly recapitulate previous *ex-vivo* work demonstrating a PARylation-dependent increase in Arc, C-Fos, and Egr1 expression following high-frequency electrical stimulation of rat cortical neurons (Visochek et al., 2016). We did not detect a PARylation-dependent increase in C-fos or Egr1; this partial discrepancy between the two studies may be due to the different experimental settings and species used (rats vs. mice).

Intra-cerebral and systemic PARP inhibitor applications were done before to manipulate reconsolidation and extinction of fear conditioning successfully (Elharrar, Dikshtein, Meninger-Mordechay, Lichtenstein, & Yadid, 2021; Goldberg et al., 2009; Inaba, Tsukagoshi, & Kida, 2015). However, the downstream effects of this inhibition were unknown. Our findings expand the role of PARylation in modulating behaviors to include stress coping. Further, we filled the gap in knowledge between the behavioral effects of PARP inhibition and the observed PARylation-dependent long-term potentiation and increased expression of some immediate-early genes after *ex-vivo* stimulation.

Our experiments were focused on several potential target genes; further research is needed to explore the transcriptome-wide effects of FST-induced PARylation and PARP inhibition. In addition, we examined the impact of FST and PARP inhibition in the cerebral cortex to closely follow the tissue selection done in previous reports (Fontán-Lozano et al., 2010; Goldberg et al., 2009; Visochek et al., 2005). Future studies are required to decipher the effects of FST on PARylation and gene expression in different subregions of the cortex and other relevant brain regions, such as the hippocampus and the amygdala. We recognize these limitations and present our findings as a first step towards understanding the PARylation-dependent mechanisms of activity-dependent gene expression regulation required for proper stress-coping behavior.

## Acknowledgments

This work was funded by an internal grant from Ariel University. The Porsolt forced swim test icon by DBCLS https://togotv.dbcls.jp/en/pics.html is licensed under CC-BY 4.0 Unported https://creativecommons.org/licenses/by/4.0/.

